# A host-pathogen interaction screen identifies *ada2* as a mediator of *Candida glabrata* defences against reactive oxygen species

**DOI:** 10.1101/248955

**Authors:** Ilias Kounatidis, Lauren Ames, Rupal Mistry, Hsueh-lui Ho, Ken Haynes, Petros Ligoxygakis

## Abstract

*Candida glabrata (C. glabrata*) forms part of the normal human gut microbiota but can cause life-threatening invasive infections in immune-compromised individuals. *C. glabrata* displays high resistance to common azole antifungals, which necessitates new treatments. In this investigation, we identified five *C. glabrata* deletion mutants (*Δada2, Δbas1*, Δhir3, *Δino2* and *Δmet31*) from a library of 196 transcription factor mutants that were unable to grow and activate an immune response in *Drosophila* larvae. This highlighted the importance of these transcription factors in *C. glabrata* infectivity. Further *ex vivo* investigation into these mutants revealed the requirement of *C. glabrata ADA2* for oxidative stress tolerance. We confirmed this observation *in vivo* whereby growth of the *C. glabrata Δada2* strain was permitted only in flies with suppressed production of reactive oxygen species (ROS). Conversely, overexpression of *ADA2* promoted *C. glabrata* replication in infected wild type larvae resulting in larval killing. We propose that *ADA2* orchestrates the response of *C. glabrata* against ROS-mediated immune defences during infection. With the need to find alternative antifungal treatment for *C. glabrata* infections, genes required for survival in the host environment, such as *ADA2*, provide promising potential targets.

## Introduction

*Candida glabrata* is a small, asexual, haploid yeast and is the second most frequent cause of candidiasis after *Candida albicans*, accounting for ~15-25% of clinical cases [1-3]. Despite its normally asymptomatic presence in the human gut microbiota, it can cause severe invasive infections in immune-compromised individuals and hospitalised patients. Such infections are associated with high mortality rates and prolonged hospital stays, consequently increasing healthcare costs. A number of risk factors including the use of central venous catheter devices and treatment with antibiotics have been associated with the development of candidiasis [2].

Although grouped as a *Candida* species, *C. glabrata* is much more closely related to the baker’s yeast *Saccharomyces cerevisiae*, a relatively non-pathogenic yeast, than to *C. albicans* [8,9]. It is proposed that *C. glabrata* and *S. cerevisiae* shared a common ancestor which underwent a whole genome duplication event [8]. Comparative genomic analysis found that these genomes lie phylogenetically distinct from the *Candida* clade containing many pathogenic yeasts, characterised by their unique translation of the CUG codon as serine rather than leucine [10, 11].

In agreement with its phylogenetic position, *C. glabrata* lacks many attributes believed to be key mediators of fungal pathogenicity in other *Candida* species such as the secretion of hydrolytic enzymes and the ability to form hyphae [4-7]. A primary virulence attribute of *C. albicans*, the leading cause of Candidiasis, is its ability to switch from a yeast to filamentous form upon certain environmental cues, which enables *C. albicans* to actively penetrate host cells. These hyphae extend filaments into the host cells, releasing hydrolytic proteases and lipases which lead to the eventual disruption of host cellular function [12]. *C. glabrata* however, is haploid and can only grow in the yeast form. Therefore, with the absence of hyphal formation, the mode of invasion of *C. glabrata* probably differ from that of *C. albicans*.

Nonetheless, *C. glabrata* is still pathogenic to humans and therefore must rely on other distinct strategies to invade and persist in infected individuals. Although the exact mode of entry is unclear the ability of *C. glabrata* to invade host tissue was demonstrated in a chicken embryo model of infection where *C. glabrata* cells were found to cross the chlorio-allantoic membrane (CAM) [12].

*S. cerevisiae* has been known to form agar invasive pseudohyphal under *in vitro* starvation conditions and it has been reported that *C. glabrata* also form pseudohyphae *in vitro* [12, 13]. *In vivo, C. glabrata* persist through endocytosis with hardly any host cell damage. Because of this low host cell damage, the cytokine profile of *C. glabrata-infected* epithelia differs significantly from that of *C. albicans-infected* cells [16]. Infection with *C. albicans* leads to a stronger pro-inflammatory cytokine response than *C. glabrata* due to the presence of hyphae and host cell damage. As a result, this leads to a strong neutrophil infiltration typical of *C. albicans* infection whereas infection with *C. glabrata* is associated with mononuclear cells [17, 18].

Once within the host, survival and the establishment of infection depends on the ability of *C. glabrata* to mount efficient responses to a changing, stressful environment acquire often limited nutrients and effectively evade the immune response.

The intrinsic resistance of *C. glabrata* to oxidative stress is of particular note [19, 20]. The oxidative burst elicited by immune cells is a first line of defence against invading microorganisms [21]. The ability to detoxify such reactive oxygen species (ROS), via the expression of detoxifying enzymes (catalase and superoxide dismutase) and production of antioxidants glutathione and thioredoxin, is important for surviving immune attack [22]. For example deletion of *SKN7*, a transcription factor mediating the oxidative stress response, attenuates *C. glabrata* virulence in a murine model of disseminated candidiasis [23]. The *C. glabrata* oxidative stress response is homologous to its *S. cerevisiae* counterpart [11] therefore additional mechanisms must contribute to increased oxidative stress resistance in *C. glabrata*. The above suggest that knowing the networks orchestrating *C. glabrata* responses to the host environment would be highly beneficial in understanding *C. glabrata* infection. Once *C. glabrata* has expanded to internal organs, its high resistance to common azole antifungals makes it hard to treat [10, 24-26]. Other high cost antifungals including echinocandins (eg. caspofungin), anidulafungin and micafungin can also induce resistance [27-29].

In light of the increased incidence of drug resistant *C. glabrata* infections, successful alternative treatment of *C. glabrata* perquisites further investigation of its pathogenicity. To this direction current mammalian models do not provide the most ideal approach to identify virulence factors because of their high cost, labour-intensive procedure, low statistical resolution and ethical concerns [30]. Nonetheless, virulence factors in *C. glabrata* have been identified in these systems, including adhesins and cell-bound proteases [31-33]. However, as large genomic libraries with deletion mutants become available in clinically relevant pathogens, invertebrates provide an alternative approach for individually investigating virulence factors and host immune responses. *Drosophila* is a small, inexpensive to rear model organism with a variety of possible genetic manipulations. Additionally, its interaction with *Candida* has been studied extensively, making it an excellent tool for screening antifungals, investigating pathogenicity and identifying novel virulent factors [30, 34-37].

The *Drosophila* Toll pathway is activated upon encountering a fungal infection in addition to Gram-positive bacterial infection [38-40]. Glucan-binding protein (GNBP3) binds to fungal β-1,3-glucan that activates a proteolytic cascade that results in cleavage of Spätzle [39, 41]. Additionally, the protease Persephone (PSH) is able to detect the activity of microbial virulence factors that also triggers the Toll pathway through the transcription factor, Dorsal-related immunity factor (DIF), which moves to the nucleus and regulates targets genes including induction of the antimicrobial peptide (AMP) gene *drosomycin (drs*) [39-41, reviewed in 42]. An additional pathway in epithelial immunity is the Imd pathway (for immune deficiency). There, a transmembrane or intracellular peptidoglycan recognition protein (PGRP-LC and PGRP-LE respectively) form a receptor-adaptor complex with IMD itself (a RIP1 homologue), which associates with FADD (the Fas-associated death domain protein) then recruiting the caspase-8 homologue DREDD (reviewed in 42]. Briefly, the signal is transmitted through the TAK1 kinase to the IκB-Kinase (IKK) on the one hand and JNK on the other. In turn, IKK (IKKα and IKKγ-otherwise known as Kenny) phosphorylates the N-terminal domain of the NF-κB homologue Relish, while DREDD cleaves the C-terminal [43]. N-terminal Rel is then free to move to the nucleus and regulate transcriptional targets including induction of antimicrobial peptide (AMP) genes [44]. In addition, Reactive Oxygen Species (ROS) are an important defence in controlling microbial invasion in the gut, regulated by the *Drosophila* dual oxidase gene (dDuox) [45].

Previous work from our laboratory involved *Drosophila* as a model organism to investigate pathogenicity of *C. albicans* [30, 37]. We determined that GI infection of wild type larvae with wild type *C. albicans* resulted in 83% of larvae reaching adulthood whereas 70% of larvae deficient for NF-κB-driven pathways developed into adults. In particular, the gut commensal bacteria community was beneficial as only 58% of germ-free larvae survived to adults while even less immune-compromised germ-free larvae (22%) developed into adults following infection with *C. albicans* [30]. Despite *C. albicans* being restricted to the gut, it still caused systemic infection as detected by *drs* expression from the fat body. Using a similar feeding protocol, we established a GI infection model for *C. glabrata* in *Drosophila*. We screened a library of deletion mutants including 196 *C. glabrata* transcription factors and we identified five that did not grow or activate systemic immunity in larvae. From this study, we propose that *ADA2* mediates *C. glabrata* defences against ROS. *C. glabrata* deficient for *ADA2* was highly sensitive to very low concentrations of H_2_O_2_ (a hallmark of oxidative stress) *ex vivo*. Inside the host, the *ADA2* mutant was only able to grow in ROS-suppressed *Drosophila* while wild type *C. glabrata* with an additional copy of *ada2* was able to grow and kill wild type larvae. We propose that *ADA2* could be a potential target to diminish *C. glabrata* growth during infection.

## Results

### The host response following *C. glabrata* GI infection

To establish an oral infection screen for a *C. glabrata* mutant library, we developed a third instar larval yeast-feeding protocol (see material and methods for details). Briefly, we fed larvae a *C. albicans* or *S. cerevisae* or *C. glabrata/banana* mixture while in parallel to each infection, larvae fed with banana only, served as our control. For these experiments we made use of third instar *w*^*1118*^ larvae (a wild type strain and the genetic background of the mutant fly strains used in this study). First, we determined the dynamics of host immune triggering after GI infection with reference yeast strains compared to the banana-only control (Figs 1A-C). As a measure of Toll pathway activity we monitored *drs* mRNA levels at specific time-points (Fig. 1A-C). On average, 24h was the time point where the most elevated *drs* gene expression levels was seen (Figs 1A-C). However, there was a significant difference in the capacity of the three fungi to activate *drs* gene expression with *C. albicans-induced* activation almost 10x more compared to *S. cerevisiae* and *C. glabrata* (Fig. 1A-C). Nevertheless, compared to non-infected larvae (Fig. 1D), there was a robust fat body induction of a *drs-GFP* marker in larvae infected with the reference *C. glabrata* strain Cg2001H (Fig. 1E). This confirmed a systemic response after feeding and was reminiscent of the same result in *C. albicans* infection [37]. In addition, there was significant increase in ROS production in Cg2001H-infected larvae over and above the banana control (Fig. 1F). Finally there was a significantly reduced survival of GI infected larvae at 24h compared to both the banana control and to non-infected larvae (Fig. 1G). Of note, compromised survival of *w*^*1118*^ was not an effect of that specific genetic background as it was also observed in *Oregon*^*R*^ and *w*^*1118*^*; drs-GFP* larvae (Figs. S1A, S1B).

**Figure 1:**
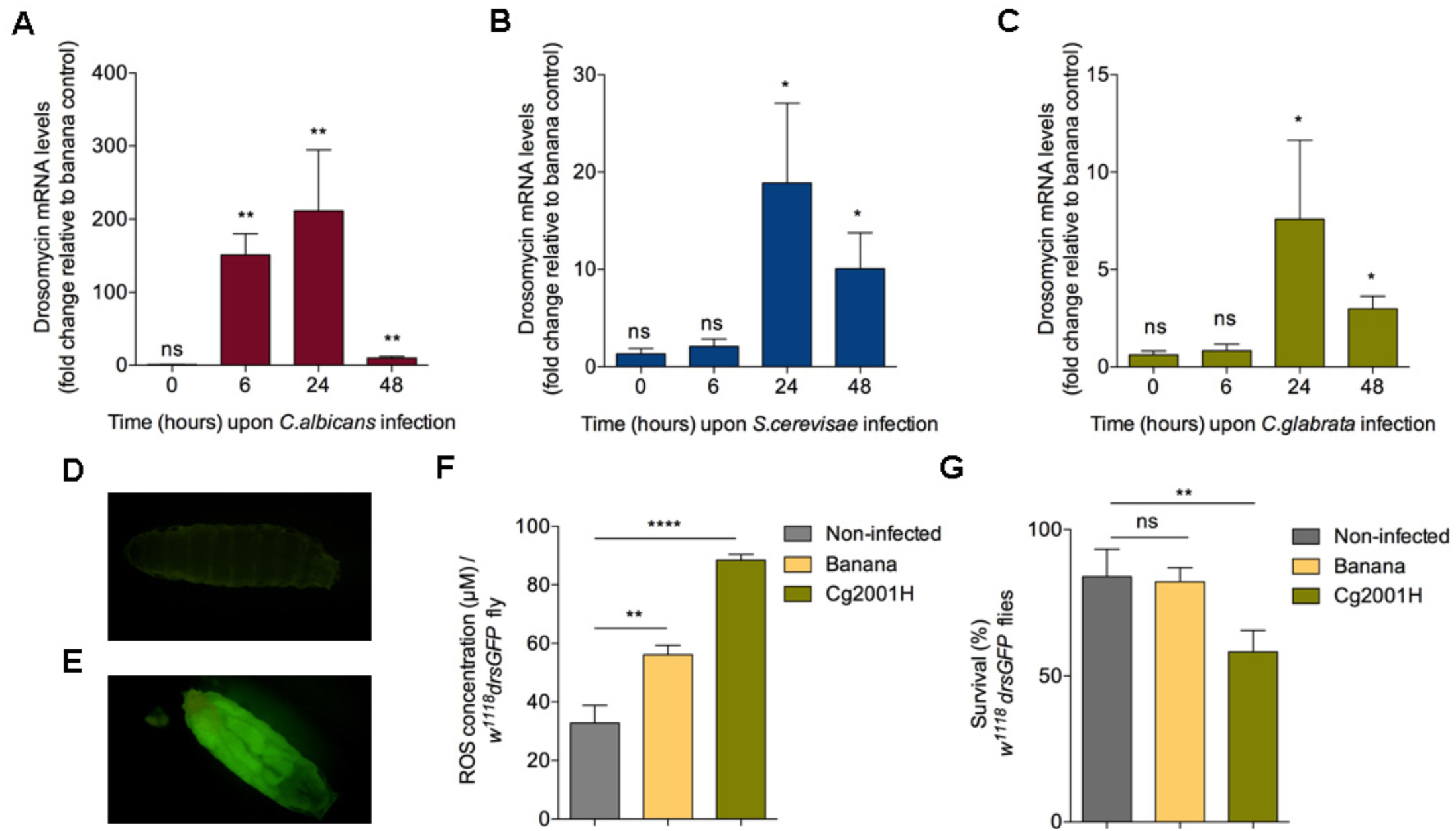
The host’s response to gastrointestinal fungal infection. Quantification of *drs* mRNA levels following oral infection of *w*^*1118*^ with wild type yeast strains at selected time points (0, 6, 24 and 48 hours). Infection with **(A)** *C. albicans*, **(B)** S. *cerevisiae* and **(C)** C. *albicans. Drosomycin* levels at each time point are relative to the banana control of the corresponding time point. Error bars represent the standard deviation of 5 separate experiments; ns p>0.05, *p<0.05, **p<0.01 indicate significant values when compared to banana control of the corresponding time point. **(D)** Non-infected *w*^*1118*^*;drsGFp* white pre-pupae visualised under a fluorescent stereoscope. **(E)** *w*^*1118*^*;drsGFp* white prepupae 48 hours post-infection with Cg2001H. **(F)** Quantification of ROS levels in noninfected, banana-fed and *C. glabrata* infected *w*^*1118*^*;drsGFp* larvae. **(G)** Percentage of larvae surviving to pupae in non-infected, banana fed and *C. glabrata* infected *w*^*1118*^*;drsGFp* larvae. Error bars represent the standard deviation of 3 independent biological experiments; ns p>0.05, *p<0.05, **p<0.01, ***p<0.001 and ****p<0.0001 indicate significant values when compared to non-infected larvae.

### Exploring individual strain infectivity in a library of *C. glabrata* deletion mutants

After establishing the feeding protocol to outline the host response to *C. glabrata* GI infection, we used the system to screen a library of *C. glabrata* deletion mutants (TFKO library) for their ability to activate the Toll pathway. The library included deletions of 196 *C. glabrata* transcription factors (TF), with well-characterised roles in other yeast species [46]. In this manner, we would be able to pinpoint to gene regulatory networks organised by specific transcription factors that are involved in infectivity. Table S1 presents all the strains used for the screen. We used *w*^*1118*^*; drs-GFP* as a proxy for immune induction and virulence. Five of the 196 *C. glabrata* TF mutants (*Δada2, Δbas1, Δhir3, Δino2, Δmet31*) did not activate *drs* expression at 24h following infection (Fig. 2A). To ascertain that this phenomenon was due to the deleted TFs, complementation strains were constructed. Complemented strains were able to activate *drs* to a degree that was statistically indistinguishable from the *C. glabrata* reference strain, *Cg2001H* (Fig. 2B). In contrast, this was not the case when the TFKO strains were complemented with just the empty vector (Fig. S2A; for a list of the complementation strains used see Table S1). We used Colony Forming Units (CFUs) to monitor the presence of *Cg2001H* inside GI infected larvae. *W*^*1118*^*; drs-GFP* larvae presented similar kinetics with *w*^*1118*^ larvae (Fig 2C and Fig. 2D respectively), with *Cg2001* cleared at approx. 64h post-feeding. In contrast, immune-deficient *w*^*1118*^*; dif-key* larvae showed delayed clearance with a considerable CFUs of *Cg2001H* still at 96h post-feeding (Fig. 2E). Direct comparison between *w*^*1118*^ and *w*^*1118*^*; dif-key* larvae showed that at 48h post-infection, there was a significant difference in CFUs with a 24h difference in clearance (Fig. 2F). Nevertheless, the five TFKO strains that did not activate *drs* in our screen were cleared significantly faster than *Cg2001H* with zero or much lower CFU count at 24h post-infection (Fig. 2G).

**Figure 2:**
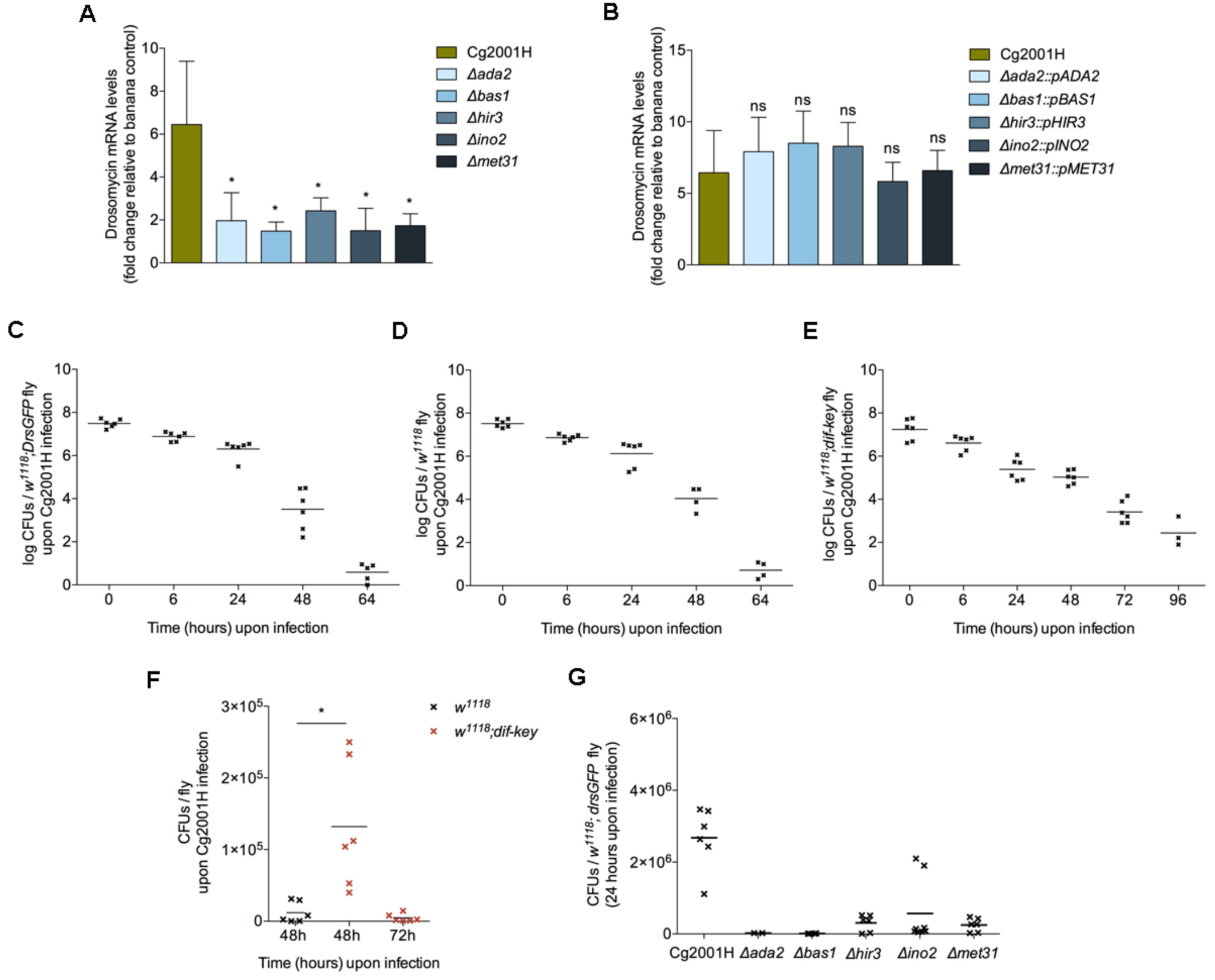
Gastrointestinal infection with deletion mutant strains. **(A)** *Drosomycin* mRNA gene expression levels relative to banana control in *w*^*1118*^*;drsGF* larvae and the TFKO deletion mutants (hit strains only), 24 hours following infection with Cg2001H. **(B)** Quantification of *drosomycin* gene expression following infection of *w*^*1118*^*;drsGFp* with Cg2001H and the corresponding complemented strains of the TFKO deletion mutants relative to banana control 24 hours post-infection. Error bars represent the standard deviation of 5 separate experiments; ns p>0.05, *p<0.05 indicate significant values when compared to *w*^*1118*^*;drsGFp* infected with Cg2001H. CFUs/fly of *C. glabrata* in **(C)** *w*^*1118*^ *;drsGFp* larvae, **(D)** *w*^*1118*^ larvae and **(E)** *w*^*1118*^ *;dif-key* larvae at selected time points post-infection with Cg2001H. **(F)** CFUs/fly of Cg2001H in *w*^*1118*^ 48 hours post-infection and in *w*^*1118*^*;dif-key* 48 and 72 hours after infection. **(G)** CFUs/fly of Cg2001H and the 5 deletion mutants (*Δada2, Δbas1, Δhir3, Δino2* and *Δmet31*) in *w*^*1118*^*;drsGFp* larvae 24 hours postinfection.

### *Ex vivo* phenotypic characterisation of *C. glabrata* TF mutants failing to activate Drosophila immunity

To pinpoint a mechanism by which the *C. glabrata* mutants may be cleared in the larval host, the mutants were tested for growth on several conditions relating to the host environment. This included nutrient limitation tolerance, pH and temperature sensitivity and susceptibility to antifungals (Fig. 3A; for a list of all *ex vivo* conditions tested see Table S2). *C. glabrata Δbas1* and *Δhir3* grew similarly to the parental strain Cg2001 under all stress conditions tested while *Δino2* was susceptible to 0.01%SDS (Fig. 3A). In addition, *Δmet31* was susceptible to H_2_O_2_, 20mM HU and 0.01% SDS (Fig. 3A). The most susceptible phenotypes were observed for *C. glabrata Δada2* which was susceptible to growth on 10 out of 13 of the tested conditions relating to growth under decreased temperatures, nutrient limitation, oxidative stress, cell membrane stress, cell wall stress and antifungal drug tolerance (Fig 3B, Table S3).

**Figure 3:**
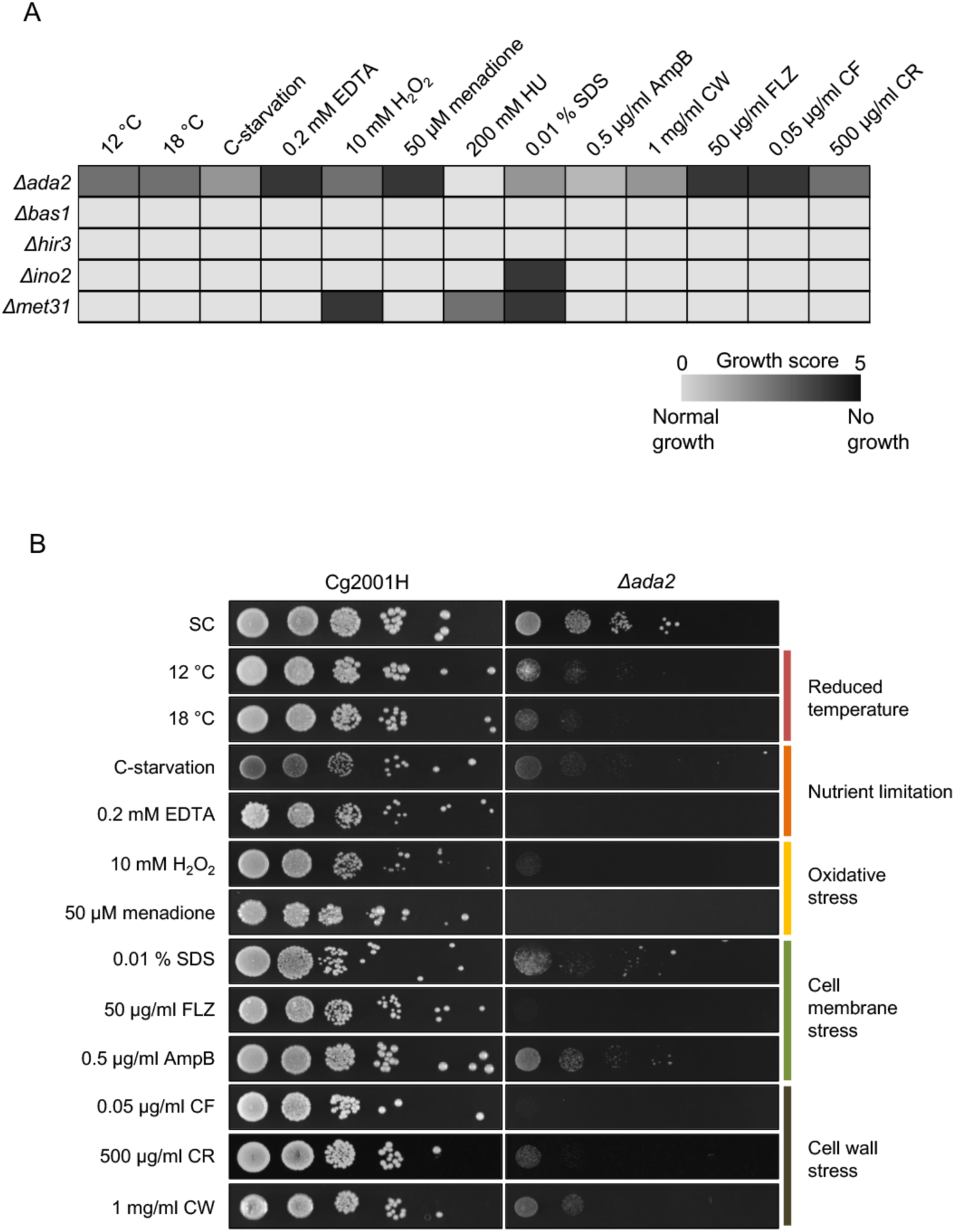
*Ex vivo* phenotypic growth of the deletion mutant strains. **(A)** Selected *C. glabrata* mutants were tested for growth under stress conditions targeting a variety of cellular processes and structures. Ten-fold serial dilutions of parental strain Cg2001H and deletion mutants Δada2, Δbas1, Δhir3, Δino2 and Δmet31 were spotted onto SC (synthetic complete) agar supplemented with the described chemical agents. Plates were incubated at 30 °C and scored for growth. Stress conditions for which all strains showed normal growth are not included. HU = hydroxyurea, SDS = sodium dodecyl sulphate, AmpB = amphotericin B, CW = calcofluor white, FLZ = fluconazole, CF = caspofungin, CR = congo red. **(B)** Serial dilution images of Δada2 phenotypes relative to parental strain Cg2001H. Stress conditions are grouped according stress type and targets.

The five TF deletion mutants were further investigated for any possible dysfunctions, which may relate to their reduced ability to persist in the larval host. Initially, we monitored growth under non-stressed conditions at 30°C. Outside the host, all mutant strains displayed growth curves comparable to the reference strain (Fig. S3A). These growth rate measurements indicated that these TF mutants were not growth-defective. However, *C. glabrata Δada2* and *Δbas1* deletion mutants displayed an increased generation time (Fig. S3B).

The susceptibility of *C. glabrata Δada2* to oxidative stress inducing agents H_2_O_2_ and menadione were of particular interest since increased levels of ROS were recorded upon *Cg2001H* infection in earlier experiments (Fig 1F). Therefore, we explored the hypothesis that *ADA2* was indispensable for resistance to toxic levels of oxidative stress, something that would be the case when encountering the localised epithelial immune response of the *Drosophila* GI tract.

### The role of the *C. glabrata* TF *ADA2* in *Drosophila* GI infection

The *Drosophila* dual oxidase (dDuox) has been shown to regulate ROS in the fly intestine [45]. When *dDuox* was knocked down in enterocytes (*w*^*1118*^*; np1-GAL4; Duox*^*RNAi*^), larval survival following GI infection with *Cg2001H* was significantly reduced (Fig. 4A). This showed that absence of ROS production increased susceptibility of larvae to GI infection. In contrast, survival of these larvae following infection by the *Δada2* deletion mutant was significantly improved compared to *Cg2001H* (Fig. 4A). This showed that when both host ROS production as well as pathogen ROS defences were absent larval survival was largely restored. This was not the case when *Δada2* was complemented with a wild type copy of the *ada2* gene (Fig. 4A). Consistent with the role of *Δada2* in the process, complementation of the *Aada2* mutant with an empty vector did not compromise larval survival (Fig. 4A). Finally, adding an extra copy of the *ada2* gene in the *Cg2001H* reference strain made the latter more virulent than normal when host ROS production was supressed (Fig. 4A). This underlined the role of *ada2* as a major regulator of *C. glabrata* virulence beyond ROS defences in the context of gastrointestinal infection. In contrast, none of the other mutants compromised larval survival when overexpressed (Fig. 4B).

**Figure 4:**
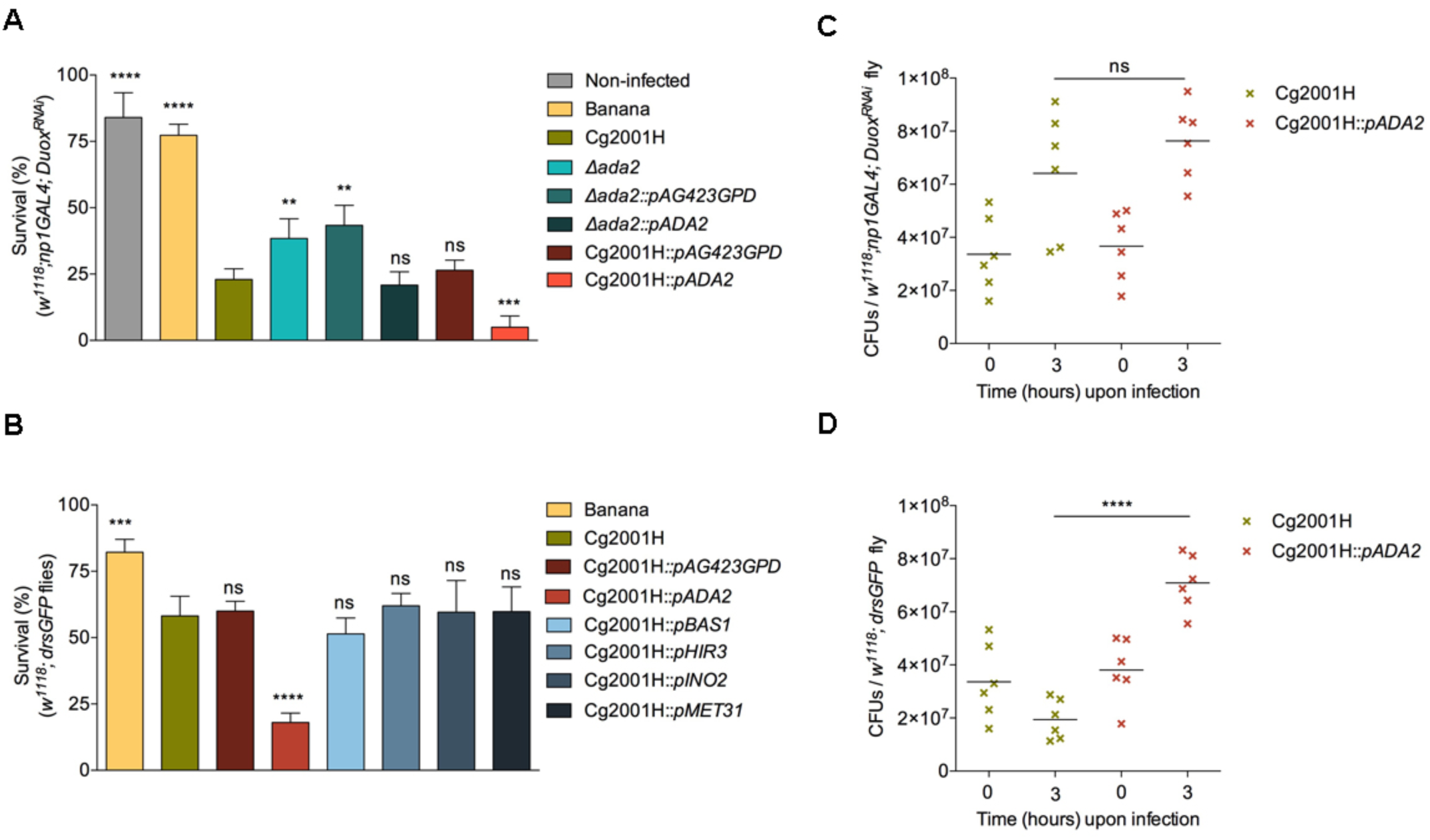
Oral infection of ROS suppressed larvae. **(A)** The percentage of ROS suppressed larvae developing into pupae in non-infected, banana fed larvae, or larvae infected by various strains including Cg2001H, *Δada2* deletion strain, *Δada2* deletion strain with empty vector (*Δada2::pAG423GPD), Δada2* complement strain (*Δada2::pADA2*), wild type *C. glabrata* with empty vector (Cg2001H::pAG423GPD) and *ada2* overexpression strain (Cg2001H::pADA2). **(B)** The percentage of *w*^*1118*^*;drsGFp* larvae developed into pupae in banana-fed larvae, infected larvae with wild type Cg2001H, *C. glabrata* with empty vector (Cg2001H::*pAG423GPD*) and *ada2*, *bas1*, *hir2*, *ino2* and *met31* overexpression strains **(C)** CFUs/fly 0 and 3 hours after infection with Cg2001H and *ada2* overexpression strain in ROS suppressed larvae (*w*^*1118*^*; np1GAL4; Duox*^*RNAi*^). **(D)** CFUs/fly 0 and 3 hours after infection with Cg2001H and *ada2* overexpression strain in *w*^*1118*^ larvae; ns p>0.05, *p<0.05, **p<0.01, ***p<0.001 and ****p<0.0001 indicate significant values when compared to larvae infected with Cg2001H.

The above increased pathogenicity was further confirmed in various strains like the GAL4 and RNAi lines used to supress ROS production (Fig S4A and S4B respectively), *w*^*1118*^ (as the genetic background of all of the mutants used; Fig. S4C) and *Oregon*^*R*^ flies (as an extra genetic background; Fig. S4D). Overexpression strains of the rest of the hits were also constructed (Table S1) and their pathogenicity was examined in the same host background like the *ada2* overexpression strain. Survival of larvae showed that hyper virulence of *ada2* overexpression was unique among the other TFKO hits of the original screen (Fig. 4C).

CFU measurements revealed that there was an expansion of both *Cg2001H* and *Cg2001H* overexpressing *ada2 (Cg2001H::pADA2*) when *Duox* gene expression was knocked-down (Fig. 4C) resulting in increased larvae lethality. However, in case of infection of a wild-type host strain (with normal ROS production) there was a significant increase in CFUS of *Cg2001H::pADA2* in contrast to the clearance of the reference *Cg2001H* strain (Fig. 2C and Fig. 4D). This meant that an extra copy of *ada2* allowed increased growth of the pathogen. Moreover, as seen by the larval survival assays, elevated growth of *Cg2001H::pADA2* compromised the host significantly more than the wild type *C. glabrata*. However, this growth advantage was lost when larvae were unable to produce ROS. Taken together with the inability of *ada2* to grow in even the lowest of H_2_O_2_ concentrations, the above results suggest that the TF *ada2* is important for regulating *C. glabrata* defences against host-generated ROS.

## Discussion

*C. glabrata* is becoming an important problem in persistent hospital infections. Starting as a commensal microbe it can generate systemic challenges that have an increasingly incurable outlook. The growing resistance of *C. glabrata* to various antifungals including azoles underlines the above and indicates the need to develop new drugs and/or therapies to target this pathogen [3, 27-29]. To this end, we screened a *C. glabrata* TFKO mutant library [46] to identify possible novel virulence factors responsible for *C. glabrata* pathogenicity at the level of the intestine. Five candidate genes were identified and their ability to infect the host was explored further. Five deletion strains showed accelerated clearance by the host gut environment. Considering that gut conditions are a new stress environment for the pathogen we followed a series of stress response tests including oxidative stress sensitivity, antifungal sensitivity, pH and temperature sensitivity. Between the five TFKO deletion mutant hits, *Δada2* deletion showed severe growing limitations under oxidative stress response tests. Therefore, this particular mutant was studied further using both ROS suppressed as well as wild type *Drosophila* strains.

### The side of the host

The synthesis of ROS (including H_2_O_2_, hydroxyl radicals and superoxide) as an immune defence is the first line immune response by phagocytes upon encountering a fungal infection in humans [45]. Similarly, the production of both ROS and AMPs are important features of *Drosophila* epithelial immunity [45, 47, 48]. ROS can cause damage to DNA, RNA, and proteins and endorse the oxidative degradation of lipids in cell membranes. GI infection of *Drosophila* larvae by *C. glabrata* increased ROS levels significantly when compared to non-infected or banana-only treated controls. When the percentage of larvae surviving to pupae was determined, we found that although feeding with banana activated a low level ROS induction (Fig. 1F) this did not affect the survival of the organism. However, infection with *Cg2001H* did impact survival with approx. 60-65% of larvae developing to pupae. Oral infection with *C. glabrata* activated epithelial immunity in the gastrointestinal tract by inducing both the ROS production mechanism as well as leading to a systemic activation of the Toll pathway. Lack of the Toll and Imd pathways just delayed the clearance of the pathogen, implying a major role for ROS in the equilibrium between host and pathogen. The crucial role of ROS was further supported by recording significantly increased levels of host lethality in larvae with ROS suppression.

### The side of the pathogen

Five deletion mutants from the TFKO library showed accelerated clearance and inability to induce an immune response inside the host. Growth curves of the 5 deletion mutants showed that all the mutants were able to successfully grow at 30°C in YPD media, outside of the host. Moreover, the generation times for the 5 deletion mutants were similar, with *Δada2* and *ΔbasJ* displaying though a marginally shorter generation time than the other strains. Nonetheless, all the deletion strains did not display any detectable growth defects that would be the cause of its rapid elimination from the host. The ability of the deletion mutants to grow under different stress conditions included tests for nutrient limitation tolerance, sensitivity to antifungals, defects in cell wall/membrane and sensitivity to oxidative stress conditions, for example. All but one strain were able to grow under all the conditions, albeit reduced and limited growth was observed. However, *Δada2* deletion mutant showed high sensitivity to the oxidative stress conditions with very restricted or no growth. As previously mentioned, wild type *C. glabrata* has shown to grow in extremely high H_2_O_2_ conditions [49]. Thus, the inability of *Δada2* mutants to grow in low H_2_O_2_ conditions indicated that the *ada2* gene is essential for its resistance to the ROS-mediated immune defense in the *Drosophila* gastrointestinal tract. To this direction, *Δada2* mutant was investigated further using ROS-suppressed flies.

### *Δada2* mutant and the interaction with host-generated ROS

The open reading frame (ORF) of *ada2* in *C. glabrata* is CAGL0K06193g and is known to be involved in the transcriptional activation of RNA polymerase II. Moreover, ortholog(s) are also implicated in chromatin binding and histone acetyltransferase. This suggests that the gene function of *ada2* is very broad and is likely to be involved in the activation of various proteins. *Ex vivo* experiments clearly showed that *Δada2* deletion mutants were highly sensitive to various oxidative stress conditions suggesting that among the many functions of this gene, it is also involved in resistance to oxidative stress like the ROS-mediated immune defence in *Drosophila*. In *C. albicans*, ROS affects only cells exposed to phagocytosis and epithelial immunity rather than those in systemic infection [50]. Mechanisms to avoid or defend against host ROS in this fungus include extracellular anti-oxidant enzymes of the super dismutase (SOD) family of enzymes [51, 52].

Larvae with supressed ROS production were able to survive to pupae like wild type strains, indicating that the lack of ROS synthesis in the gut had no impact on survival under normal conditions. However, infection with wild type *C. glabrata* resulted in a significant decrease in survival. Interestingly, the percentage of ROS suppressed larvae developing into pupae was significantly more when infected with *Δada2* than when challenged with wild-type *C. glabrata*, Cg2001H. This indicated that in the absence of the *ada2* gene, ROS suppressed larvae were able to survive better. This made *ada2* a good candidate to mediate transcription related to ROS resistance in *C. glabrata*. When *ada2* gene was re-introduced, virulence restored. Moreover, when this gene was overexpressed, survival of ROS suppressed flies decreased significantly. This confirmed that the *ada2* gene is necessary to survive the ROS-mediated immune defence of the host. The fungal load increased after three hours postinfection of *ada2* overexpression strain in ROS suppressed flies. Moreover, in wild type (*w*^*1118*^) larvae with functioning Duox, *ada2* overexpression strain was able to persist and expand three hours after infection while the fungal load of wild type *C. glabrata* decreased. This indicates that an additional copy of the *ada2* gene makes this strain hypervirulent, significantly reducing the survival of the host and enabling it to expand within the host. Contrastingly, survival of *w*^*1118*^*;drsGFP* larvae to pupae when infected with the remaining 4 overexpression strains of the deletion mutant hits (*Δbas1, Δhir3, Δino2* and *Δmet31*) was similar to wild type *C. glabrata* highlighting the non-random effect of *ada2* overexpression.

The induction of ROS has been shown to be a rapid and an important immune defense following natural infection in *Drosophila* [45]. Flies that lack the ability to generate ROS upon natural infection succumb to infection. Ha and colleagues demonstrated ubiquitous expression of *Duox-RNAi* resulted in increased mortality following natural infection with *Ecc15*. Likewise, when *Duox-RNAi* was restricted to the gut, flies displayed a similar level of mortality. Interestingly, when *Duox-RNAi* was introduced to the main immune tissues in systemic immunity in *Drosophila* (the fat body/hemocytes), survival of flies was unaffected [46]. This indicates that ROS plays a major role in the host resistance during natural infection.

A previous study by Schwarzmüller and colleagues in 2014 [46] where the original deletion library was constructed analysed the sensitivity of the deletion strains under few basic conditions and their resistance to fungicides. A total of 14 deletion strains displayed marked hypersensitivities to azoles such as fluconazole and voriconazole. *Ada2* deletion strain was included in this group and its above sensitivity was further confirmed by additional fungicides test conducted in our research. Thus, a potential drug component that could target specifically *ada2* silencing, could be prove very effective in efforts of eliminating *C. glabrata* pathogen.

## Materials and Methods

### Drosophila stocks and genetics

The following stocks were used: *w*^*1118*^ (BL #6326), *Drs-GFP* [53], *dif-key* [54], OregonR. Mutant strains were isogenised by backcrossing 10 times in *w*^*1118*^ background, thus refer in *w*^*1118*^ the manuscript as *w*^*1118*^*; drs-GFP* and *w*^*1118*^*; dif-key*.

### Infection Experiments

We followed a modified gastrointestinal infection model [44] based on a previously established bacterial gastrointestinal model [37]. In more details, prior to infection, 10mL of yeast extract peptone dextrose (YPD) (Y1375, Sigma-Aldrich, USA) was inoculated with a single colony of yeast and incubated on a rotating shaker at 30°C at 120 rpm for 16 hours. The inoculum was pelleted by centrifuging at 4°C at 4000 rpm for 4 minutes and the supernatant was discarded. The pellet was washed in 10mL of 1X PBS and centrifuged again at 4°C at 4000 rpm for 4 minutes and the supernatant was discarded. A single banana was homogenised and 450μL was added to 2mL eppendorf tubes. To this, 250μL of pathogen (OD=200) was also added. As a control, banana without pathogen was used. Five-day (third instar) larvae were collected by washing from the fly food and caught in a sieve. They were starved for one hour before infection. ~100 larvae were added to each eppendorf tube, plugged with breathable foam with space for fermentation and allowed to feed for 30 minutes. The mixture then was transferred to a standard fly medium and incubated at 30°C. Lethality of larvae following infection was determined by counting surviving pupae. 48 hours after infection.

### TFKOs mutants screen

The transcription factor knockout (TFKO) library screen was conducted following the above oral infection protocol in *w*^*1118*^*; drs-GFP* 5-day old larvae. Two days after infection, pupae were visualised under UV light to confirm that they were unable to activate immunity. Further to this, *drs-GFP* were infected with the complement strain of each hit to re-confirm that when the knocked-out gene was re-inserted, immunity would be activated.

### Yeast counts from larvae by counting CFUs

Following infection, larvae were washed in 100% ethanol, rinsed in sterile water and transferred to normal fly food. Larvae were homogenised (at indicated time points) in 200μL YPD media (Y1375, Sigma-Aldrich, USA). Serial dilutions were made and 50μL, plated on YPD plates (Y1500, Sigma-Aldrich, USA). Plates were incubated at 30o and colonies were measured accordingly.

### Microscopy

Larvae were visualised on a GFP stereo dissecting microscope (Leica MZFIII, UK), and images captured using KyLink software (v2.0, Japan).

### Gene expression analysis

Drosomycin expression was determined from five independent biological samples consisting of 5 infected third instar larvae following infection. Samples were compared to banana-fed flies of the corresponding time point. Larvae were collected at the desired time points and washed in 100% ethanol and sterile water. RNA was extracted using Purification Plus Kit (48400, Norgen – Biotek, Canada) and cDNA was prepared from 0.5 μg total RNA using Maxima First Strand cDNA Syntesis Kit (K1672, Thermo Scientific, UK). Triplicate cDNA samples were amplified with the SensiFASAT SYBR No-ROX Kit (BIO-98020, Bioline, UK) in a Corbet Rotor-Gene 6000 QPCR machine (Qiagen, UK) according to the manufacturer's protocols. Expression values were calculated using the DDCt method and normalized to rp49 expression levels [56]. Primers Used: rp49(forw): AAGAAGCGCACCAAGCACTT CATC, rp49(rev): TCTGTTGTCGATACCCTTGGGCTT, drosomycin(forw): AGTACTTGTTCGCCCTCTTCGCTG, drosomycin(rev): CCTTGTATCTTCCGGACAGGCAGT.

### Measuring production of reactive oxygen species (ROS)

The amount of ROS was measured by quantifying the amount of hydrogen peroxide produced per larvae immediately after oral infection. The samples included three independent biological samples of 10 third instar larvae from non-infected, banana-fed and infected with C. glabrata samples. The amount of hydrogen peroxide produced from the sample was determined using Amplex^®^ red hydrogen peroxide/peroxidase assay kit (A22188, Invitrogen, USA) following manufacturer’s instructions.

### C. glabrata culture conditions

C. glabrata strains were cultured in YPD (1% yeast extract, 2 % bacteriological peptone, 2 % glucose) at 30 °C. A final concentration of 200 μg/ml nourseothricin (Werner BioAgents) was added for selection of C. glabrata deletion mutants. Complementation and overexpression strains were cultured in SC broth (0.69 % yeast nitrogen base without amino acids, 2 % glucose) supplemented with CSM single drop out (-His) mixture (Formedium). YPD and SC plates contained 2 % agar.

### C. glabrata deletion mutant construction

Construction of C. glabrata transcription factor deletion mutants used in this study was previously described [46].

### C. glabrata complementation and overexpression strain construction

C. glabrata ORFs previously cloned into the pDONR221 entry vector using GATEWAY cloning technology [57] were shuttled into a pAG423GPD-ccdB destination vector (AddGene) using LR clonase. Destination vectors carrying the C. glabrata ORF were transformed by electroporation into the corresponding C. glabrata deletion mutant for complementation or into the parental C. glabrata Δhis3 strain for overexpression. Empty pAG423GPD-ccdB vector was transformed into C. glabrata as a control. Correct transformants were selected for growth on SC ‐his media. Three independent transformants of each strain were collected.

### Phenotypic screening

Overnight cultures of C. glabrata strains were normalised to OD600 0.1 in sterile water. The normalised suspensions were aliquoted into a 96-well plate and diluted ten-fold across six wells to create serial dilutions. Using a multichannel pipette, 5 μl aliquots of the serial dilutions were spotted onto SC agar plates supplemented with stress-inducing chemical agents (Supplementary table). Plates were incubated at 30 °C, unless otherwise stated, and imaged daily. Upon visual inspection, phenotypes for each mutant were scored relative to the parental strain into six categories: mild sensitivity (MS), sensitive (S), severe sensitivity (SS), no growth (NG), improved growth (IG) or no phenotype. Three biological replicates were performed for all phenotypic screens.

### Growth analysis

Overnight cultures of C. glabrata were normalised to OD600 0.1 in fresh YPD and distributed into a flat-bottomed 96-well plate in 100 μl aliquots. Optical density readings were taken every 10 minutes in a VersaMax™ Absorbance Microplate Reader set to 30 °C with shaking between reads. Growth rate was measured by calculating the doubling time for each mutant during exponential phase (OD600 0.4 – 0.8).

## Acknowledgements

We would like to thank the members of the Ligoxygakis and Haynes labs for discussions and critical reading and Peter Burns for help with fly breeding. This work was funded by the Wellcome Trust (to KH) and NC3Rs (to PL and RM). IK was supported by a postdoctoral fellowship from the Bodossaki Foundation.

